# Arabidopsis Apoplastic Fluid Contains sRNA- and Circular RNA-Protein Complexes that Are Located Outside Extracellular Vesicles

**DOI:** 10.1101/2021.10.02.462881

**Authors:** Hana Zand Karimi, Patricia Baldrich, Brian D. Rutter, Lucía Borniego, Kamil K. Zajt, Blake C. Meyers, Roger W. Innes

## Abstract

Previously, we have shown that apoplastic wash fluid purified from Arabidopsis leaves contains small RNAs (sRNAs). To investigate whether these sRNAs are encapsulated inside extracellular vesicles (EVs), we treated EVs isolated from Arabidopsis leaves with the protease trypsin and RNase A, which should degrade RNAs located outside EVs but not those located inside. These analyses revealed that apoplastic RNAs are mostly located outside EVs and are associated with proteins. Further analyses of these extracellular RNAs (exRNAs) revealed that they comprise both sRNAs and long non-coding RNAs (lncRNAs), including circular RNAs (circRNAs). We also found that exRNAs are highly enriched in the post-transcriptional modification N^6^-methyladenine (m^6^A). Consistent with this, we identified a putative m^6^A-binding protein in apoplastic wash fluid, GLYCINE-RICH RNA-BINDING PROTEIN 7 (GRP7), as well as the small RNA-binding protein ARGONAUTE2 (AGO2). These two proteins co-immunoprecipitated with each other, and with lncRNAs, including circRNAs. Mutation of *GRP7* or *AGO2* caused changes in both the sRNA and lncRNA content of apoplastic wash fluid, suggesting that these proteins contribute to the secretion and/or stabilization of exRNAs. We propose that these extravesicular RNAs mediate host-induced gene silencing, rather than RNA inside EVs.

**One-sentence summary:** The apoplast of Arabidopsis leaves contains diverse small and long-noncoding RNAs, including circular RNAs, that are bound to protein complexes and are located outside extracellular vesicles.

## INTRODUCTION

The apoplast is the extracellular space outside the plasma membrane of plant cells that comprises the cell wall, xylem, and space between cells (Steudle, 1980; Guerra-Guimarães et al., 2016). Apoplastic fluid contains water, sugars, amino acids, cell wall modifying enzymes, growth regulators, and diverse stress-related proteins (Guerra-Guimarães et al., 2016; Huber and O’Day, 2017; Narula et al., 2020; Wang and Dean, 2020; Wang et al., 2020). Recently, we and others have shown that apoplastic fluid also contains extracellular vesicles (EVs) that carry defense related proteins and small RNAs (sRNAs) (Rutter and Innes, 2017; Cai et al., 2018a; Baldrich et al., 2019; He et al., 2021). The role of EVs in plant-microbe interactions is thus an active area of investigation.

It has been shown that sRNAs from both plants and pathogens can hijack microbe or host RNA interference pathways to induce trans-kingdom gene silencing (Weiberg et al., 2013; Niu et al., 2015; Wang et al., 2017; Hou et al., 2019; Huang et al., 2019; Schaefer et al., 2020). Expression of sRNAs in plants that target pathogen genes has been used to confer resistance to diverse fungal, nematode and insect species (Nowara et al., 2010; Koch et al., 2013; Mamta et al., 2016; Qi et al., 2019). However, it is not clear how sRNAs are transferred between plant and pathogen cells. To avoid degradation, it is speculated that extracellular RNAs (exRNAs) need to be either tightly associated with RNA-binding proteins or encapsulated within EVs (Rutter and Innes, 2017; Koch and Wassenegger, 2021), but whether EVs and/or RNA-binding proteins are required for RNA secretion or movement within the apoplast is still under investigation.

Previously, we have reported that apoplastic wash fluid contains diverse species of small RNAs, including microRNAs (miRNAs), small interfering RNAs (siRNAs), and a previously overlooked class of tiny RNAs (tyRNAs; 10 to 17 nt) with unknown functions (Baldrich et al., 2019). In that study, we showed that apoplastic tyRNAs co-purified with EVs when using a density gradient. Notably, siRNAs and miRNAs were largely missing from density gradient-purified EVs, although they were present in total apoplastic wash fluid. These observations suggested that EVs may not be the primary carrier of apoplastic siRNAs and miRNAs (Baldrich et al., 2019). In support of this hypothesis, analysis of apoplastic siRNAs derived from transgenic expression of a hairpin RNA in Arabidopsis revealed that >70% of these were located outside EVs (Schlemmer et al., 2021).

Although density gradient centrifugation is a preferred method for obtaining highly pure EV preparations (Rutter and Innes, 2020), it is still possible for large RNA-protein complexes to co-purify with EVs, or RNAs to adhere to the surface of EVs, thus most work published to date, including our own, has not established whether plant EV-associated RNA is located inside or outside EVs. To eliminate extravesicular RNA-protein complexes and RNA attached to the surface of EVs, it is necessary to treat purified EVs first with proteases to remove any RNA-binding proteins and then with RNase to degrade the released RNAs (Rutter and Innes, 2020).

Recently, He et al. (2021) identified several RNA-binding proteins in the apoplast of Arabidopsis leaves that might be responsible for loading sRNAs into EVs, including ARGONAUTE1 (AGO1), ANNEXIN1 and 2 (ANN1 and ANN2), and RNA HELICASE11 and 37 (RH11 and RH37). Protease protection assays indicated that these proteins are all located inside EVs. However, this work did not include a protease plus RNase treatment, thus did not distinguish between sRNAs located outside EVs in RNA-protein complexes versus sRNAs located inside EVs (He et al., 2021). Similarly, Cai et al. (2018), used micrococcal nuclease treatment to show that sRNAs that had co-pelleted with EVs were protected from degradation. However, the lack of prior protease treatment likely left RNA-protein complexes intact, thus this analysis also did not distinguish between sRNAs located in RNA-protein complexes versus those located inside EVs.

Although plant EVs have only been reported to contain sRNAs and tyRNAs, mammalian EVs have been reported to carry sRNAs as well as lncRNAs, including circular RNAs (circRNAs) (Xu et al., 2020b). circRNAs are covalently closed, single-stranded circles derived from back-splicing reactions of RNA Polymerase II transcripts, whereby a splice donor site at the 3’ end of an exon fuses to a splice acceptor site at the 5’ end of the same exon, or another upstream exon (Fu and Ares, 2014; Wang et al., 2021). circRNAs have been shown to play a regulatory role in multiple biological processes, including immune responses in both mammalian and plant systems (Hu et al., 2019; Mahmoudi et al., 2019; Fan et al., 2020; Zhang et al., 2020b). One mechanism by which circRNAs are thought to regulate gene expression is through acting as sponges for both miRNAs and RNA-binding proteins, and thereby sequestering them. Such sequestration can impact RNA transcription, splicing, and translation (Hansen et al., 2013; Jeck and Sharpless, 2014; Bose and Ain, 2018; Panda, 2018). Fan et al. (2020) demonstrated that circRNAs from rice are involved in immune responses to the fungal pathogen *Magnaporthe oryzae*. Several circRNAs in rice leaves were detected only upon infection with *M. oryzae*. Furthermore, this work showed that overexpression of one specific circRNA enhanced rice immunity to *M. oryzae* (Fan et al., 2020), indicating that circRNAs may represent an important component of plant immune systems. However, whether circRNAs are secreted by plant cells, as they are by mammalian cells, has not yet been reported.

To understand the possible function of exRNAs in plants, we analyzed the sRNA and circRNA content of Arabidopsis apoplastic fluid both inside and outside EVs, as well as the RNA-binding proteins associated with these RNAs. Our data reveal that apoplastic fluid contains diverse RNA species, including sRNAs and lncRNAs (100 >1,000 nt), many of which appear to be circRNAs. The great majority of both sRNAs and lncRNAs were found to be located outside EVs. However, this extravesicular RNA is protected against degradation by RNases via association with RNA-binding proteins. The presence of abundant extravesicular sRNA- and circRNA-protein complexes in the apoplast suggests that these RNAs may play a central role in plant-microbe interactions and also contribute to host-induced gene silencing.

## RESULTS

### The Majority of Apoplastic Small RNAs Are Located Outside EVs

Our previous analyses of sRNAs associated with density gradient-purified EVs revealed that EVs contain relatively few RNAs in the 21, 22, and 24 nucleotide size range, and instead are highly enriched in RNAs 10-17 nucleotides in length, so-called tiny RNAs (tyRNAs) (Baldrich et al., 2019). Those analyses, however, did not assess whether these tyRNAs were located inside or outside the EVs and did not include any apoplastic sRNAs that pelleted at 40,000g but that did not co-purify with EVs in the density gradient. To assess whether apoplastic fluid contains RNA-associated particles other than EVs, we generated sRNA libraries from pellets obtained after centrifuging apoplastic wash fluid at 40,000g for one hour (P40 pellets; see Materials and Methods). P40 pellets contain a mixture of particles, including EVs. To distinguish between RNA located inside EVs from RNA located outside EVs, we treated P40 pellets with trypsin plus RNase A, which should eliminate RNA associated with proteins located outside EVs, while leaving RNAs located inside EVs intact. As controls, we treated pellets with just the buffer or with RNase A alone. The latter should degrade free RNA but not RNA bound to proteins or located inside EVs. Separate sRNA-seq libraries were generated from each of three biological replicates of each treatment (nine libraries in total) and sequenced using an Illumina NextSeq platform. We observed that the distribution of read lengths was consistent between replicates, but substantially different between treatments (Figure 1). Control samples displayed predominant peaks at 21, 22, and 31 nt, while RNase A alone treated samples displayed peaks at 16 and 17 nt and trypsin plus RNase A treated samples displayed peaks at 10 and 12 nt. These results are consistent with our previous analyses of density gradient-purified EVs in that EVs appear to contain very few 21, 22, or 24 nt sRNAs but are enriched in tyRNAs. Significantly, these results reveal that the apoplast contains large amounts of 21 and 22 nt RNAs that are located outside EVs and bound to proteins.

**Figure 1.**
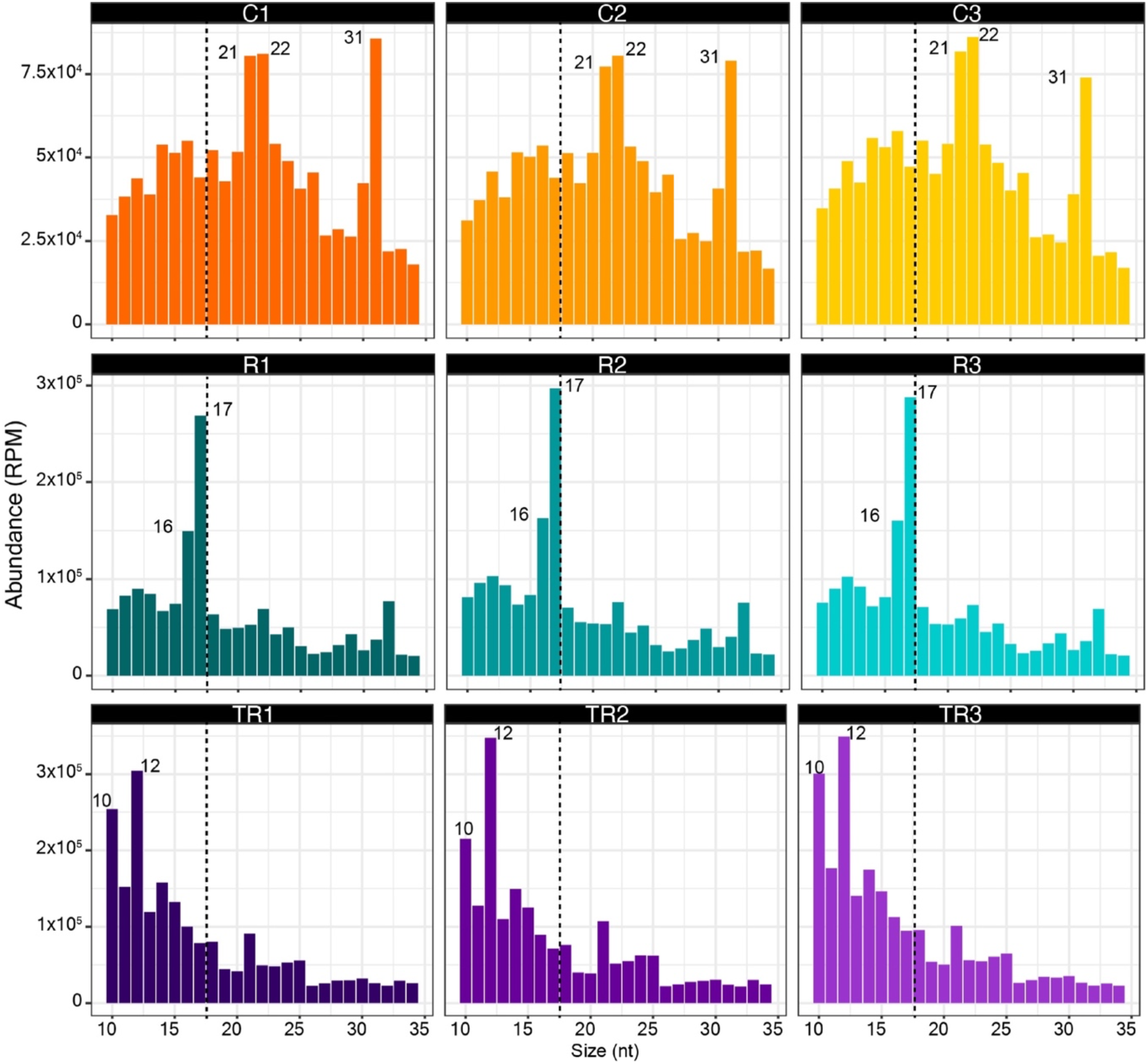
The Majority of Apoplastic Small RNAs are Located Outside EVs. Size distribution of P40 sRNAs mapping to the genome; the abundance of each size class was calculated for each P40 treatment (control (C1-C3), RNase A only (R1-R3), and trypsin plus RNase A(TR1-TR3)). The x axis indicates the sRNA size and the y axis indicates its abundance in reads per million (RPM). Data from three independent biological replicates are shown. Note the loss of 21 and 22 nt reads following treatment with trypsin plus RNase A, which indicates this size class is mostly found outside EVs.

To further understand the nature of apoplastic sRNAs and tyRNAs, we analyzed their origin. We observed that most of the sRNA reads originated from rRNAs, mRNA, and products dependent on RNA Polymerase IV (Pol IV) (Figure 2). We also observed that the treatment with RNase A and trypsin had a different impact on each RNA category. While relative representation of mRNA and rRNA categories remained fairly constant after different treatments, the representation of Pol IV-, miRNA-, small nuclear RNA (snRNA)- and transposable element (TE)-derived sRNAs increased after RNase treatment and decreased after trypsin plus RNase A treatment (Figure 2A). This pattern suggests that Pol IV-, miRNA, snRNA- and TE-derived sRNAs are mostly located outside EVs but are protected from degradation due to association with proteins. In contrast, the relative amount of tRNA-derived sRNAs decreased after RNase treatment but increased after trypsin plus RNase treatment, suggesting that tRNA-derived sRNAs are present in the apoplast as unprotected RNAs outside EVs, as well as inside EVs. In the case of tyRNAS, we observed an increase in all categories after trypsin plus RNase treatment (Figure 2B). These patterns support our previous conclusion that tyRNAs are highly enriched inside EVs (Baldrich et al., 2019).

**Figure 2.**
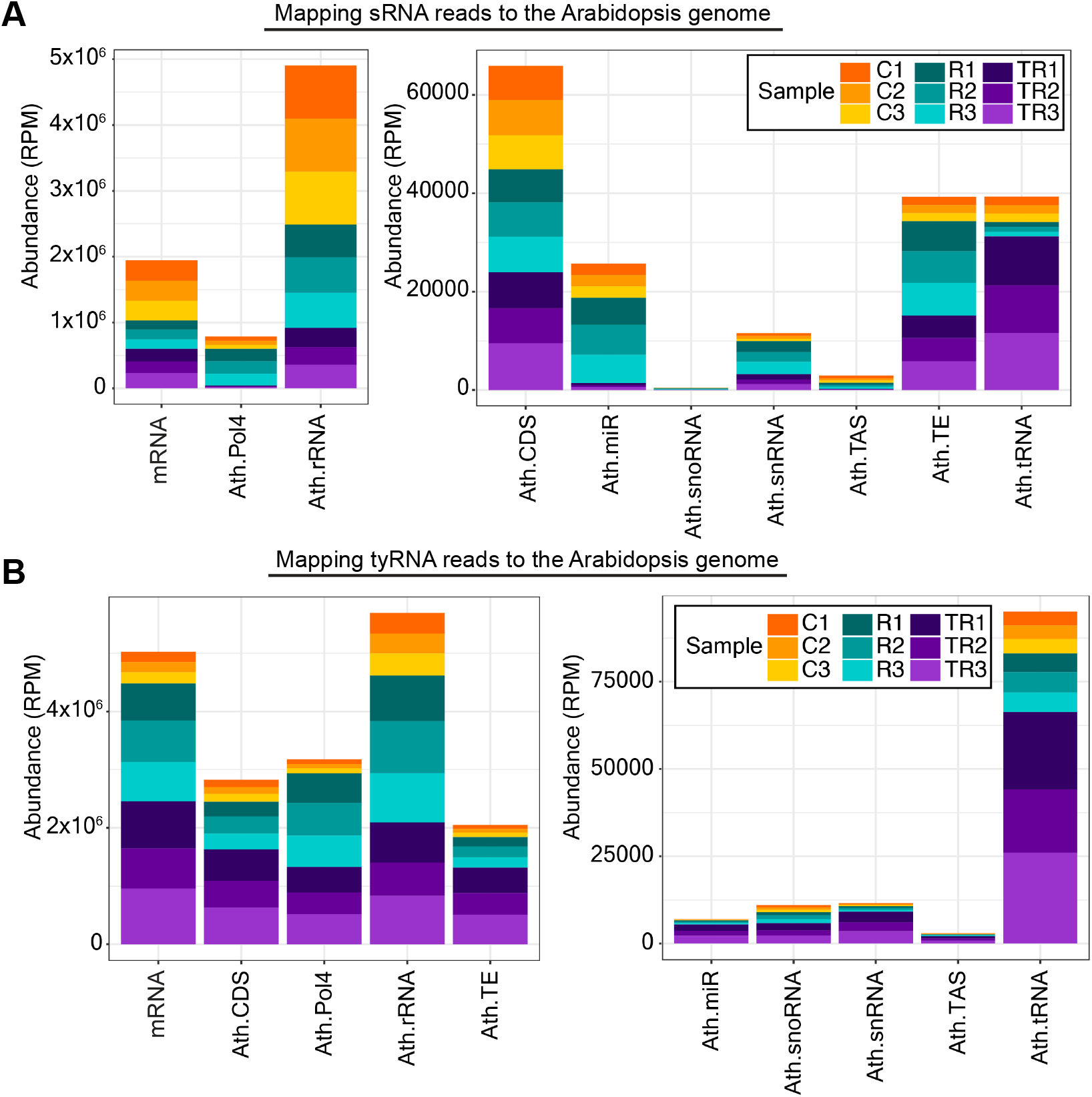
Apoplastic sRNAs Are Derived from Diverse Sources. **(A)** Specific subclasses of sRNAs are protected by proteins. sRNAs that mapped to the genome were categorized by origin and plotted by relative abundance in reads per million (RPM). Treatment with RNase A alone (R1-R3) increased the relative proportion of Pol IV-, miRNA-, snRNA- and TE-derived sRNAs, while treatment with trypsin plus RNase A (TR1-TR3) decreased their proportion, indicating that the majority of these sRNAs are protected by protein and are located outside EVs. TE, transposable element-derived RNA; snoRNA, small nucleolar RNA; snRNA, small nuclear RNA; TAS, trans-acting siRNA. **(B)** tyRNAs are mostly located inside EVs. tyRNAs that mapped to the genome were categorized by origin and plotted by relative abundance. All categories of tyRNAs increased in relative abundance upon treatment with trypsin plus RNase, indicating that they are protected against trypsin plus RNase treatment, hence are mostly located inside EVs. For both panels, the x axis indicates the RNA source, and the y axis indicates its abundance in reads per million (RPM). Data from three independent biological replicates are stacked together in a single bar plot and color coded as shown in the legend.

We further analyzed these sRNA-seq data by plotting the read-length distributions according to their origins (Supplemental Figure 1). As expected, read lengths for miRNAs and trans-acting siRNAs (tasiRNAs) displayed sharp peaks at 21 nt. Notably, this size distribution was not altered by treatment with RNase A alone, while treatment with trypsin plus RNase A eliminated the 21 nt peaks, leaving a peak at 10-12 nt. These observations further support our conclusion that sRNAs are primarily located outside EVs and are protected by RNA binding proteins, while tyRNAs are located inside EVs.

Supplemental Figure 1 also revealed that the peak at 31 nt observed in Figure 1 was almost entirely due to transcripts that overlap known Pol-IV-dependent 24 nt siRNAs (Zhou et al., 2018). Notably, this peak was eliminated by treatment with RNase A alone, leaving a peak at 16-17 nt. This observation suggests that these Pol IV-dependent transcripts are also located outside EVs but are only partially protected by RNA binding proteins. The observation that these transcripts are mostly 31 nt rather than 24 nt suggests that they are derived from precursor RNAs that did not complete maturation into 24 nt siRNAs by DICERLIKE 3 (DCL3) (Blevins et al., 2015).

### A Small Subset of miRNAs Are Enriched Inside EVs

Although we observed that apoplastic miRNAs, overall, were much more abundant outside than inside EVs, this observation did not rule out the possibility that some miRNAs might be specifically loaded into EVs and thus could be enriched inside EVs relative to the general apoplastic miRNA population. To test this hypothesis, we compared the frequencies of individual miRNAs in each sample using a differential gene expression tool (see Methods). To avoid false negatives due to low expression, we selected only miRNAs with more than one read per million (RPM) in at least one sample. This filter reduced the dataset from 427 mature miRNAs to 94. From these, 62 miRNAs displayed differential accumulation in at least one of the comparisons (Figure 3). Based on the differential accumulation pattern, we placed the miRNAs into six clades. Clade I comprised seven miRNAs that were highly accumulated in the trypsin plus RNase A treated samples compared to control and RNase A alone treated samples but were not differentially accumulated in RNase A alone treated versus control samples. This is the pattern expected for miRNAs located inside EVs, which should be protected against RNase A degradation regardless of trypsin treatment. Clade II comprised ten miRNAs that were significantly more abundant in the RNase A alone treated samples relative to the control samples, and in the trypsin plus RNase A treated samples relative to controls, with three of these also being significantly more abundant in the trypsin plus RNase A samples versus the RNase A alone samples. This pattern would be expected for miRNAs that are located both inside EVs and outside EVs, with the latter being protected against RNase digestion by proteins. Clades III and VI contained 24 miRNAs that exhibited low accumulation in trypsin plus RNase A treated samples compared to control and RNase A alone treated samples but high accumulation in RNase A alone treated samples compared to the controls. This pattern is expected for miRNAs that are located outside EVs and protected by RNA-binding proteins. The miRNAs found in Clade IV (13 total) and Clade V (8 total) exhibited low abundance in RNase A alone treated samples versus controls as well as trypsin plus RNase A alone versus controls. These are most likely miRNAs that are located outside EVs and are not protected by proteins. In summary, these data indicate that most plant miRNA species in the apoplast are located outside EVs, with only seven miRNAs apparently enriched inside EVs.

**Figure 3.**
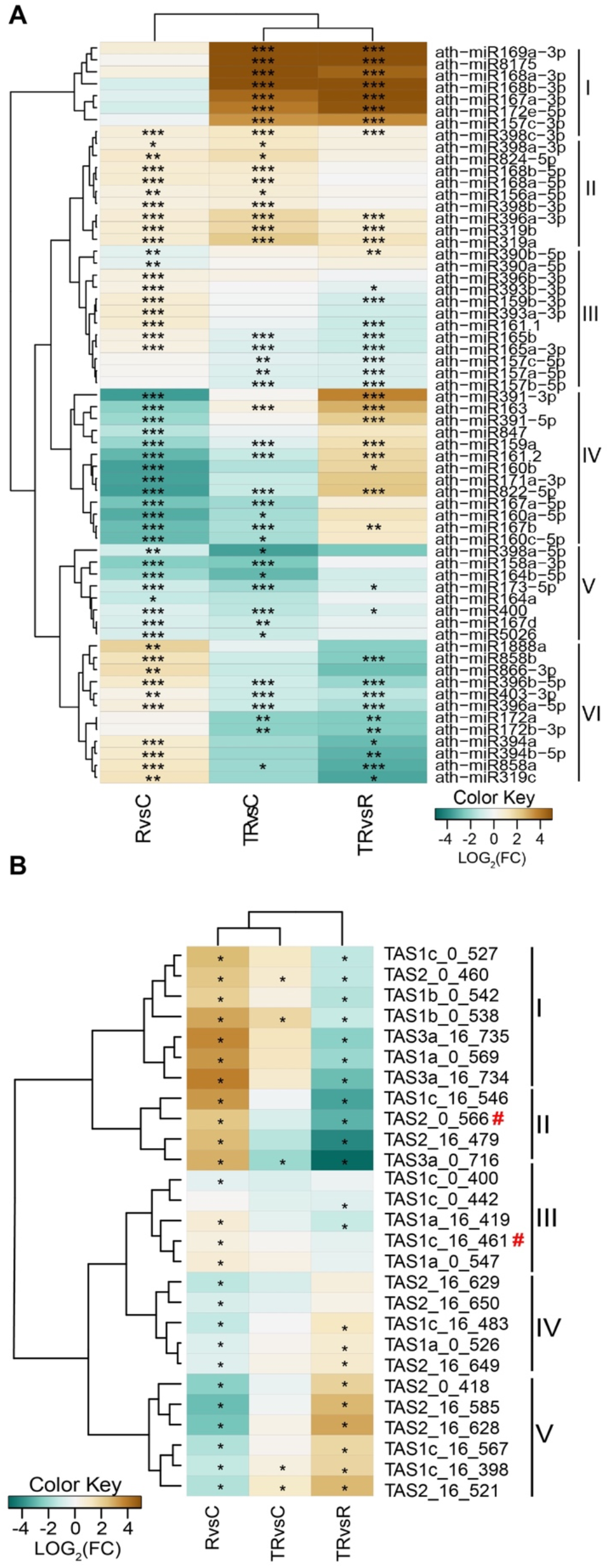
Apoplastic sRNAs are Mostly Found Outside EVs. **(A)** Apoplastic miRNAs with a minimum abundance of one read per million in at least one treatment and differentially accumulated in at least one comparison were grouped into six clades based on their relative abundance following three different treatments: RNase A alone (R), trypsin plus RNase A (TR), or no treatment (C). Heat map indicates enrichment (brown) or depletion (teal) in one treatment compared to another. **(B)** Apoplastic tasiRNAs with a minimum abundance of five reads per million in at least one treatment and differentially accumulated in at least one comparison were grouped into five clades based on their relative abundance. Red hashtags indicate tasiRNAs previously reported to mediate silencing of genes in the fungus *Botrytis cinerea* (Cai et al., 2018; He et al., 2021). Heat map indicates enrichment (brown) or depletion (teal) in one treatment compared to another.

### Apoplastic tasiRNAs Are Mostly Located Outside EVs

Trans-acting siRNAs (tasiRNAs) are a subclass of sRNAs that have been proposed to mediate interkingdom RNA interference, possibly by transfer inside of plant EVs (Cai et al., 2018a; He et al., 2021). The analyses presented in Figure 1 and Supplemental Figure 1, however, indicate that siRNAs are mostly located outside EVs. To determine whether there may be a specific subset of tasiRNAs that are preferentially loaded inside EVs, we performed a differential accumulation analysis of tasiRNAs, just as described above for miRNAs. To avoid false positives, we established a minimum cut-off of five RPM in at least one sample, reducing the number from 1581 to 27 tasiRNAs. Of these, all exhibited a differential accumulation that was statistically significant in at least one comparison (Figure 3B). Based on differential abundance in the three samples, we could group these 27 tasiRNAs into five clades.

Clade I (seven tasiRNAs) showed significantly higher relative abundance in RNase A alone treated samples compared to control samples, suggesting these tasiRNAs are located outside EVs and are protected by proteins. Consistent with this conclusion, these seven tasiRNAs were relatively less abundant in trypsin plus RNase A treated samples compared to RNase A alone treated samples. Clade II tasiRNAs (four tasiRNAs) showed a very similar pattern to that of Clade I tasiRNAs, thus are also likely to be located outside EVs and protected by proteins.

Clades III (three tasiRNAs) and IV (eight tasiRNAs) showed a relative abundance pattern opposite to that of clades I and II, with treatment with RNase A alone leading to a decrease in relative abundance compared to control samples, and treatment with trypsin plus RNase A causing an increase compared to RNase A alone treated samples. This pattern suggests that tasiRNAs belonging to clades III and IV are located outside EVs and are not protected by proteins, although why trypsin plus RNase A treatment leads to less efficient removal than RNase A alone is unclear. We speculate that residual trypsin activity in the former may be lead to a slight reduction in RNase activity.

Lastly, the five tasiRNAs included in clade V showed a pattern more similar to clades I and II, suggesting that these are located outside EVs and are mostly protected by proteins. Notably, none of the tasiRNAs showed a pattern that would be consistent with protection inside EVs, which should show a relative increase in abundance across all three comparisons.

It has been previously reported that two tasiRNAs from Arabidopsis, Tas1c-siR483 (here named Tas1c_16_461) and Tas2-siR453 (here named as Tas2_0_566) are transferred into fungal cells via extracellular vesicles. However, in our study, we found that these two TAS-derived siRNAs are present outside EVs, in association with RNA-binding proteins (indicated by red # symbol in Figure 3B).

### Apoplastic Wash Fluid Contains Long RNAs That Are Protected by RNA-Binding Proteins

The above analyses revealed that Arabidopsis apoplastic fluid contains sRNA-protein complexes that are located outside EVs, thus defining a new class of extracellular RNA in plants. Recent work in mammalian systems has revealed that mammalian cells secrete lncRNAs independent of EVs (Lasda and Parker, 2016; Preußer et al., 2018). We thus investigated whether plants might also secrete lncRNAs that are extravesicular. For these analyses, we collected apoplastic wash fluid from Arabidopsis leaves using the same protocol as used for sRNA isolation (Rutter and Innes, 2017). The resulting apoplastic wash fluid was filtered and centrifuged successively at 10,000g, 40,000g, and 100,000g. RNAs were isolated after each centrifugation step and analyzed by polyacrylamide gel electrophoresis, followed by staining with SYBR® Gold to detect nucleic acids. These analyses revealed that apoplastic wash fluid contains abundant long RNAs ranging in size from 35 nt to at least 1,000 nt (Figure 4). Most of the apoplastic RNAs were pelleted by centrifuging the wash fluid at 40,000g (P40) (Figure 4A), which we have previously shown pellets EVs (Rutter and Innes, 2017).

**Figure 4.**
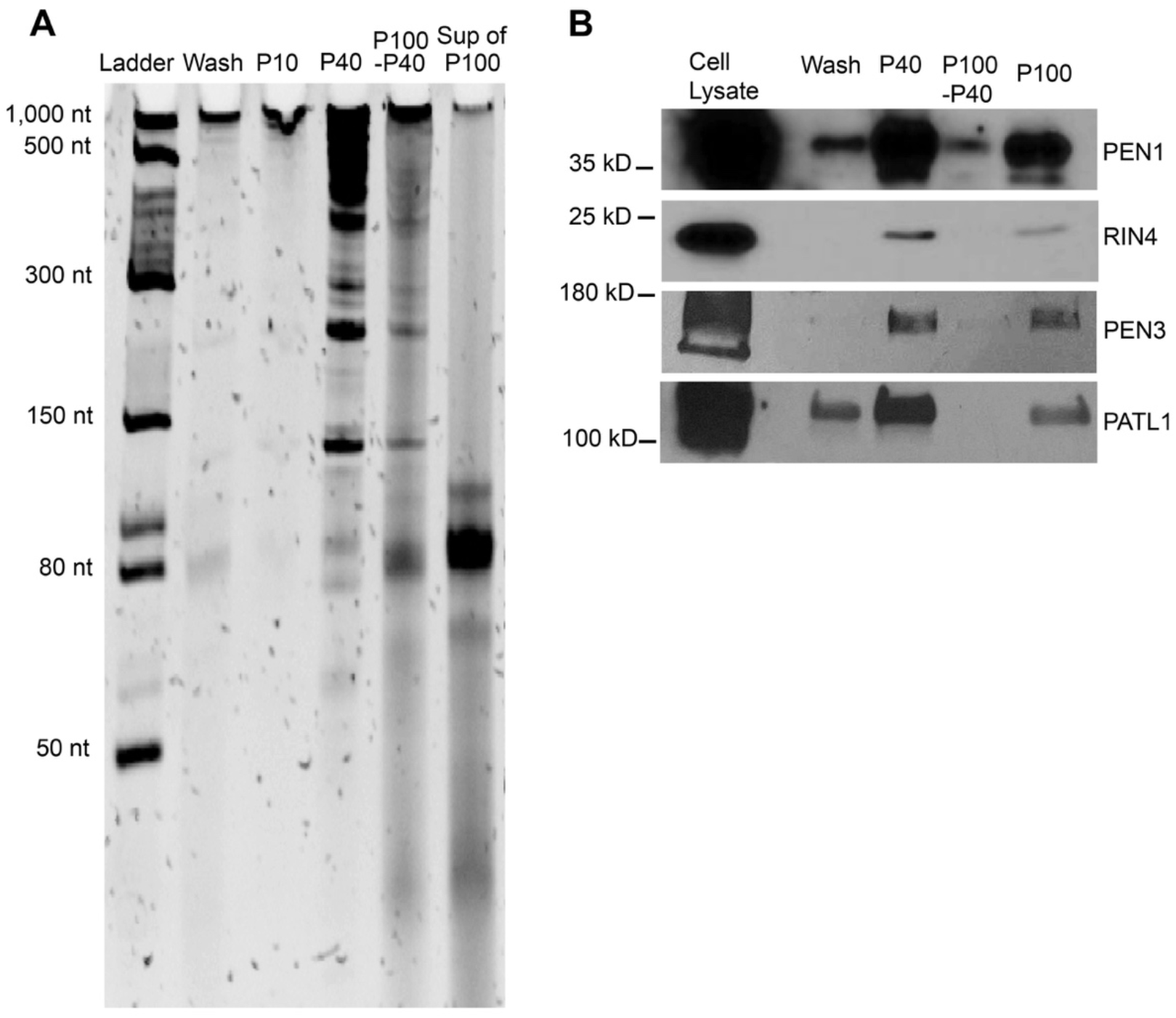
Apoplastic Fluid Contains Long RNAs. **(A)** Long RNAs are present in apoplastic fluids and can be pelleted by ultracentrifugation. RNA was isolated from the indicated fractions using TRIzol extraction and then separated on a 40% denaturing polyacrylamide gel, followed by staining with SYBR® Gold nucleic acid stain. P10, P40 and P100-40 indicate RNA isolated from pellets obtained in successive centrifugation steps at 10,000g (P10), 40,000g (P40), and 100,000g (P100-P40). ‘Sup of P100’ indicates the RNA remaining in the supernatant after the 100,000g centrifugation step. Note that the majority of the RNA larger than 50 nt is pelleted at 40,000g, indicating it is associated with particles of some kind. **(B)** EVs are pelleted at 40,000g. EV marker proteins PEN1, RIN4, PEN3, and PATL1 all pelleted at 40,000g (P40), with very little remaining in the P100-P40 pellet. ‘P100’ indicates a sample obtained by skipping the 40,000g step, going directly to a 100,000g centrifugation step following the 10,000g centrifugation step.

Some apoplastic RNAs remained in the supernatant after the 40,000g step, but were pelleted after centrifuging at 100,000g (P100-P40), which indicates the presence of some apoplastic RNAs that are not associated with EVs (Figure 4A). To assess the presence of EVs in both the P40 and P100-P40 fractions, we tested for the known EV protein markers PEN1, PEN3, PATL1, and RIN4 (Figure 4B). Consistent with our previous work (Rutter and Innes, 2017), these markers were found almost entirely in the P40 fraction. Thus, EVs are concentrated in the P40 fraction, while non-EV components, including some apoplastic RNAs, can be found in the P100-P40 fraction.

We also observed an abundant RNA smear running between 80 nt and approximately 100 nt in the supernatant following the 100,000g step (Figure 4A). This size distribution is similar to that of eukaryotic tRNAs (76-90 nt), and our previous sRNA-seq analyses on the supernatant of P40 pellets revealed abundant tRNA sequences (Baldrich et al., 2019). We thus speculate that this smear corresponds to free tRNAs.

These data indicate that apoplastic wash fluid contains multiple species of RNA, including many RNAs that are longer than 100 nt. These long RNAs must be associated with some kind of particle, as they all pellet when centrifuged at 100,000g for one hour. To distinguish RNAs encapsulated in EVs from those located in non-vesicular protein-based particles, we treated the P40 fraction with trypsin to digest extravesicular proteins and then with RNase A to digest RNAs (Figure 5). Notably, the majority of the P40 RNA was not digested by treatment with RNase A alone, but was completely degraded by treatment with trypsin followed by RNase A. To rule out the possibility that trypsin plus RNase A treatment was disrupting the integrity of EVs, we also analyzed these samples for the presence of the known EV cargo proteins PEN1, PATL1 and RIN4 (Figure 5B). All three proteins were intact in the trypsin plus RNase A treated sample but were missing from the detergent plus trypsin plus RNase A treated sample, indicating that the EVs remained intact during the trypsin plus RNase A treatment. Collectively, these results show that the majority of apoplastic RNA is located outside EVs but is protected against RNase digestion by RNA-binding proteins.

**Figure 5.**
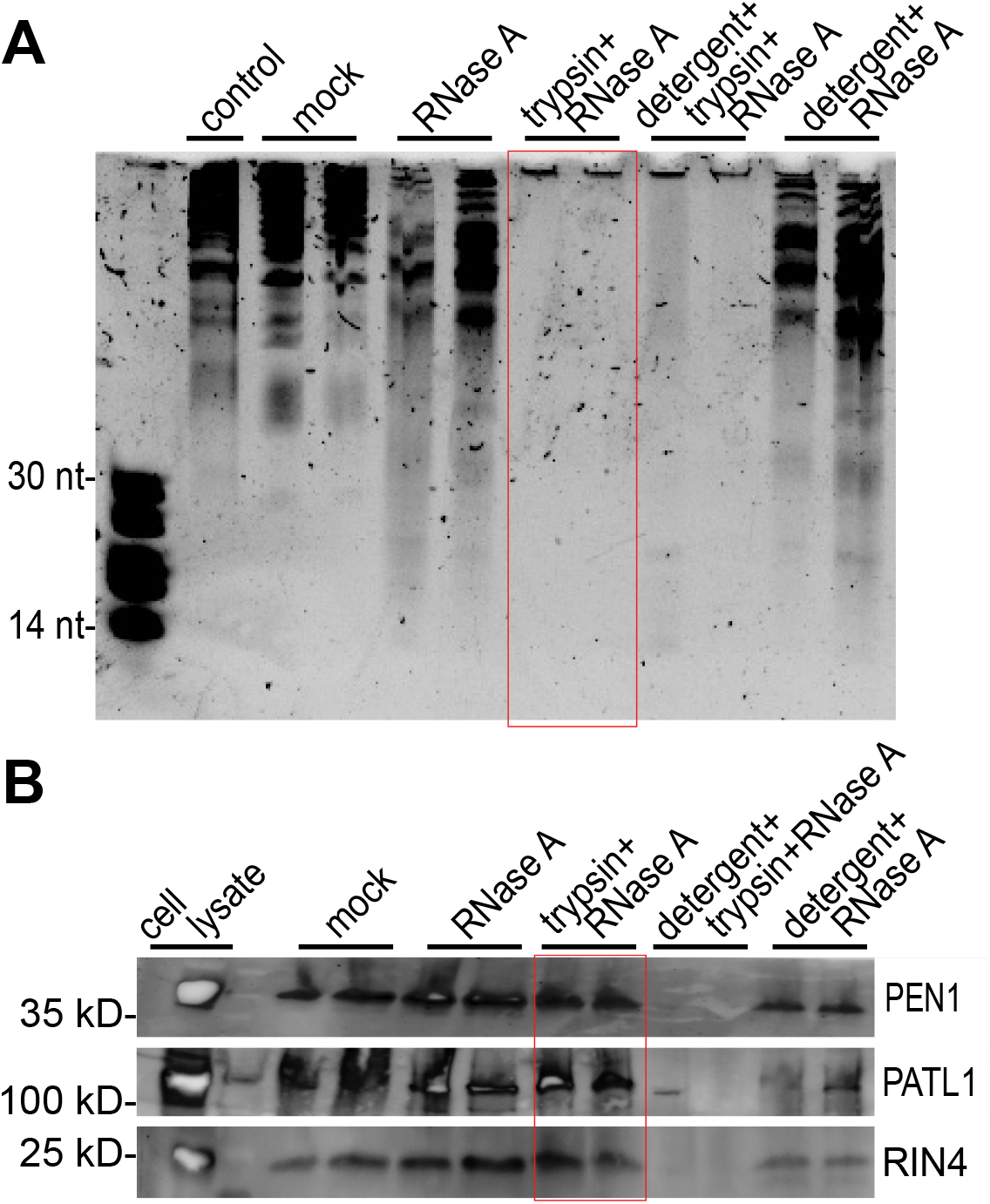
Apoplastic Long RNAs are Protected Against RNase A Digestion by Proteins. **(A)** RNase A treatment alone does not degrade most apoplastic RNA. P40 pellets were subjected to treatment with RNase A, trypsin plus RNase A, Triton X-100 detergent plus trypsin plus RNase A, or detergent plus RNase A. Control is input RNA without any treatments and kept on ice; Mock is the same RNA subjected to the same incubations as the treated RNA, but without detergent, RNase A or trypsin. RNA was analyzed using a denaturing PAGE gel as described in Figure 4. **(B)** RNase A and trypsin do not disrupt EVs. The P40 pellets from panel A were analyzed by immunoblots prior to RNA extraction to assess whether EVs remained intact. EV cargo proteins PEN1, PATL1 and RIN4 were degraded by trypsin only when detergent was included, which indicates EVs remained intact following treatment with trypsin plus RNase A.

### Apoplastic RNA Contains Circular RNAs

To our knowledge, long extracellular RNAs have not been reported previously in plants. In mammals, however, exRNAs have been extensively characterized due, in part, to their potential use as non-invasive markers for diseases such as cancer (Zhan et al., 2018). Notably, mammalian exRNAs are highly enriched in circRNAs, possibly due to their resistance to digestion by extracellular RNases (Li et al., 2015; Chen and Huang, 2018; Seimiya et al., 2020). To assess whether plant exRNAs also contain circRNAs, we performed RNase R treatment on exRNAs isolated from P100 pellets. This enzyme is a 3’ to 5’ exoribonuclease that digests most linear RNAs, including structured RNAs such as rRNA, but leaves circRNAs intact (Vincent and Deutscher, 2006). The RNase R treated RNA was then analyzed by denaturing polyacrylamide gel electrophoresis, which revealed that a large amount of RNA larger than 300 nt remained undigested, along with several distinct bands shorter than 300 nt (Figure 6). As a control, we homogenized whole Arabidopsis leaf tissue and subjected it to purification using our EV isolation protocol. The RNA obtained from this preparation displayed a banding pattern entirely different from that of the P100 RNA, and RNase R treatment eliminated all visible RNA larger than 150 nt. These results indicate that plant exRNA is enriched in circRNAs.

**Figure 6.**
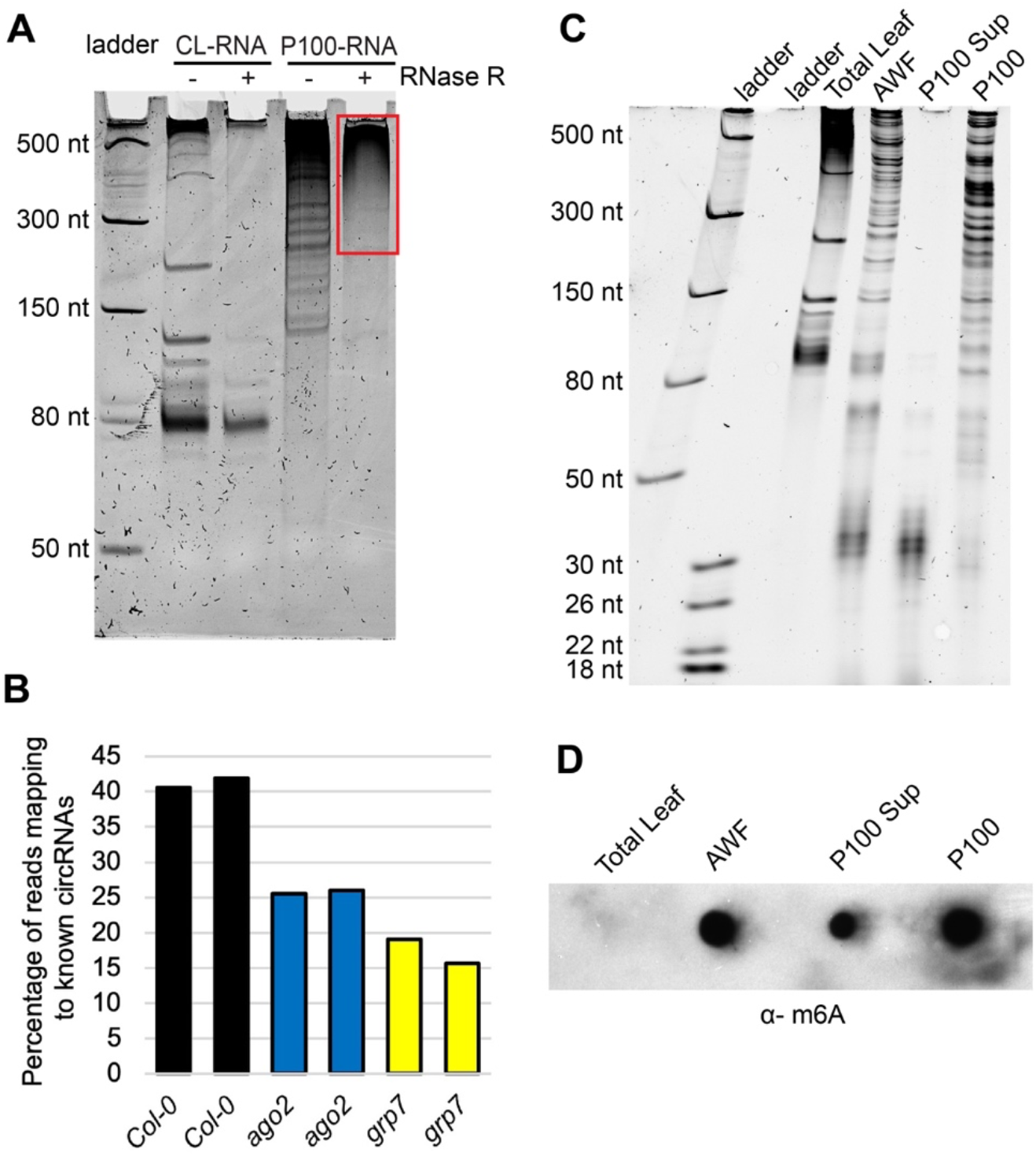
Apoplastic RNAs Are Enriched in circRNAs and m^6^A Modification. **(A)** Apoplastic fluid contains circRNAs. RNA from a P100 pellet and from total cell lysate (CL) purified using our P100 protocol was treated with RNase R, which degrades linear RNAs. RNAs were then analyzed using denaturing polyacrylamide gel electrophoresis and staining with SYBR® Gold. Red box indicates RNase R-resistant RNA. **(B)** Apoplastic RNA contains diverse circRNAs. P100 RNA was treated with RNase R to remove linear RNA, and then analyzed by RNA-seq using an Illumina NextSeq platform. Graphs indicate the percentage of reads that mapped to known Arabidopsis circRNAs for RNA isolated from wild-type, *ago2* mutant, and *grp7* mutant Arabidopsis plants. **(C)** Confirmation of RNA concentrations and integrity prior to m^6^A analysis. 200 ng of the indicated RNAs were separated on a 15% denaturing PAGE gel and stained with SYBR® Gold. **(D)** Apoplastic RNAs are enriched in m^6^A modification. 200 ng of each of the indicated RNAs from panel C were dot-blotted onto a nitrocellulose membrane and then probed with an anti-m6A antibody.

To confirm this conclusion, we generated RNA-seq libraries from P100 RNA that had been treated with RNase R and then mapped the reads from these libraries to a collection of previously identified Arabidopsis circRNAs (Chu et al., 2017), which are defined by the presence of junction fragments derived from back-splicing events (Ye et al., 2019). Consistent with our RNase R analysis, we found that apoplastic RNA contains abundant circRNAs, with over 40% of the reads mapping to known Arabidopsis circRNAs (Figure 6B).

### Apoplastic RNA is Enriched in m6A Modification

In mammalian systems, circRNA biogenesis often involves post-transcriptional modification with N^6^-methyladenine (m^6^A), which promotes back-splicing, with the resulting circRNAs containing multiple m^6^A sites (Di Timoteo et al., 2020; Zhang et al., 2020a). We thus assessed whether apoplastic RNA might be enriched in m^6^A modification. We isolated RNA from whole leaves, from total apoplastic wash fluid, from P100 pellets and from the supernatant of P100 pellets. The concentrations of these RNA preparations were then determined using a NanoDrop spectrophotometer and their concentrations equalized. To confirm that RNA samples contained equivalent amounts of RNA, 200 ng aliquots were analyzed on a PAGE gel stained with SYBR® Gold (Figure 6C). RNA samples (200 ng each) were then dot-blotted onto a nitrocellulose membrane and probed with an anti-m^6^A antibody. This analysis revealed that exRNA is highly enriched in m^6^A modification relative to total cellular RNA (Figure 6D). Notably, RNAs isolated from the P100 and supernatant of the P100 both displayed a strong signal. The latter contains mostly RNAs that are smaller than 50 nt, while the former contains RNAs larger than 50 nt. This observation suggests that both small RNAs and long RNAs may be enriched in m^6^A modification.

### Apoplastic RNA is Enriched in Intergenic RNAs

To determine the sources of apoplastic RNA, we performed Illumina-based RNA-seq analysis on RNA isolated from P40 pellets. We generated two sets of RNA-seq libraries, one using a poly(A) enrichment protocol, and one using an rRNA depletion protocol (see Methods). Analysis of the poly(A)-enriched library revealed that it contained very few products with inserts (Supplemental Figure 2), indicating that apoplastic RNA contains very little intact mRNA. This finding also indicates that there was little to no contamination with RNA from broken cells. In contrast, the second library looked as expected, thus was analyzed using Illumina sequencing. Mapping of the resulting reads to the Arabidopsis genome revealed that the majority of the reads were derived from ribosomal RNA and intergenic regions but also included a large number of reads derived from protein-coding genes (Figure 7). Notably, the latter reads included a large number of reads derived from introns, similar in number to those derived from exons, suggesting that exRNAs are enriched in incompletely spliced, or alternatively spliced RNAs. This observation is consistent with the presence of circRNAs, which often include introns.

**Figure 7.**
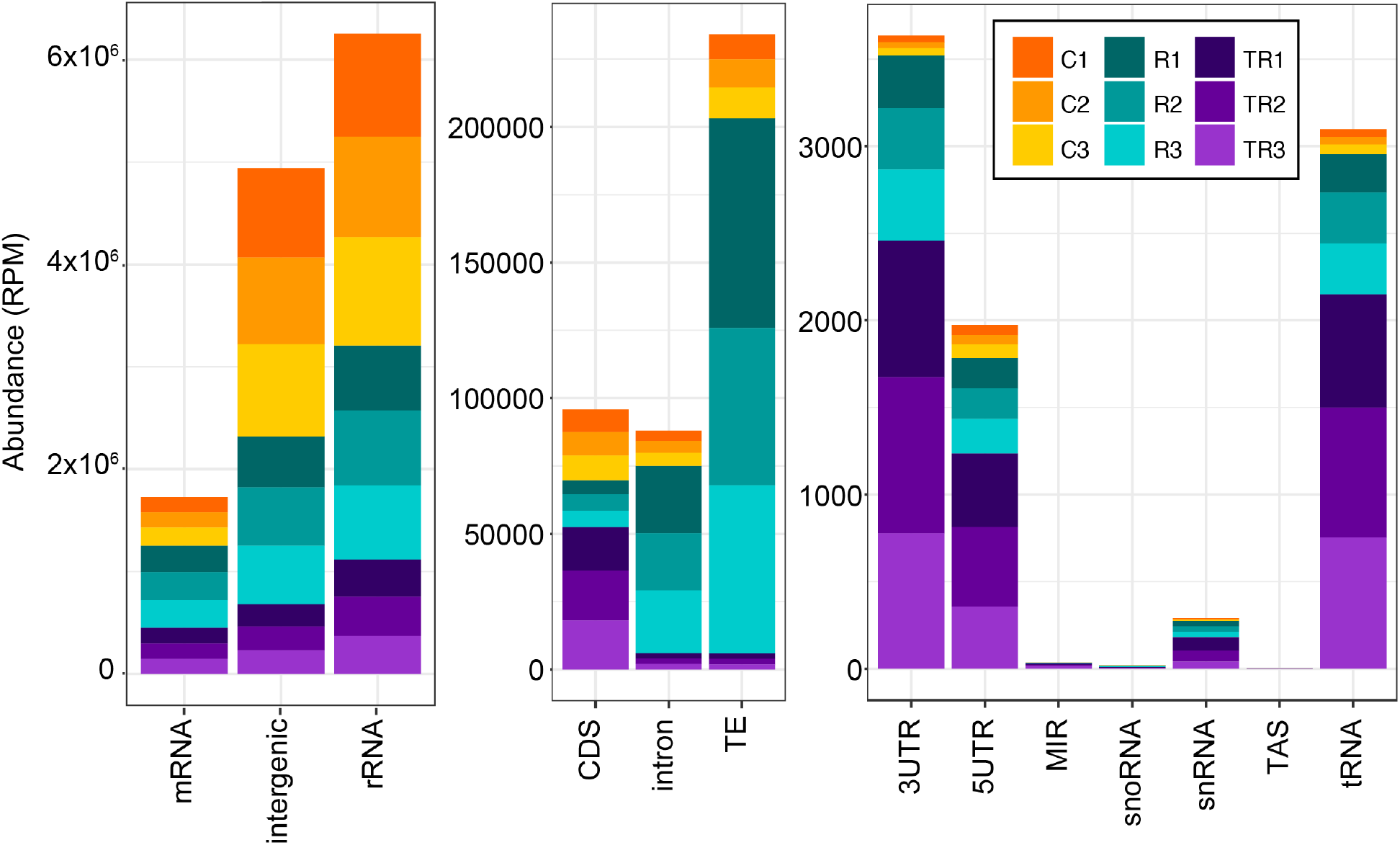
Apoplastic RNA is Derived from Multiple Sources, and Is Enriched in Intergenic RNA. RNA-seq reads were mapped to the Arabidopsis genome and categorized as indicated on the X-axis and quantified on the Y-axis by reads per million (RPM). Note the difference in scales for the three graphs, which were used to better visualize the lower abundance categories. Reads that mapped to protein coding genes (mRNA) (left graph) were further broken down into 5’ untranslated region (5UTR), 3’ untranslated region (3UTR), protein coding sequence (CDS) and intron. MIR, miRNA encoding gene; snoRNA, small nucleolar RNA; snRNA, small nuclear RNA; TAS, trans-acting siRNA-producing loci; TE, transposable elements; tRNA, transfer RNA. The values for three independent biological replicates from each of three treatments are shown. Treatments were control untreated RNA (C1-C3), RNase A-treated RNA (R1-R3) and trypsin plus RNase A-treated RNA (TR1-TR3).

To assess whether specific RNA species were associated with protein or were encapsulated inside EVs, we also made libraries from P40 pellets that were treated with RNase A alone (this should eliminate RNA that is not protected by proteins or EVs) or treated with trypsin plus RNase A (this should leave mostly RNA encapsulated in EVs). Analysis of these libraries revealed that trypsin plus RNase A reduced the relative proportion of most classes of exRNA (Figure 7), consistent with our conclusion that the vast majority of exRNA is located outside EVs but is protected by proteins. Notably, treatment with RNase A alone increased the relative frequency of RNAs that mapped to transposable elements and introns, which suggests that these RNAs are especially well-protected by proteins. In contrast, RNA reads mapping to 5’ UTRs, 3’ UTRs, and tRNAs became relatively more abundant following trypsin plus RNase A treatment (Figure 7), suggesting that these RNAs might be protected inside EVs. We interpret these data with caution, however, as these reads made up a very small fraction of the total reads. It is worth noting, also, that based on paired-end sequence reads, most of the tRNA sequences were derived from tRNA fragments and not full-length tRNAs.

### RNA-binding Proteins GRP7 and AGO2 Are Secreted into the Apoplast Independent of EVs

The above analyses revealed that apoplastic wash fluid contains abundant RNA species, including both sRNAs and long RNAs, that are protected from RNase degradation by proteins. This raised the question of what RNA-binding proteins are present in the apoplast. In our previous proteomic analyses of density-gradient purified EVs, we had identified the RNA-binding protein GLYCINE-RICH PROTEIN 7 (GRP7) as co-purifying with EVs (Rutter and Innes, 2017). GRP7 has two RNA-binding domains and binds to multiple species of RNA, including sRNAs, pre-miRNA, precursors of miRNAs and pre-mRNAs (Koster et al., 2017, Streitner et al., 2012, Nicaise et al., 2013). Arabidopsis GRP7 has been shown to participate in plant responses to pathogen infection (Fu et al., 2007, Lee et al., 2012, Nicaise et al., 2013). In addition, it is targeted by the bacterial type III-secreted effector HopU1, which blocks the interaction between GRP7 and GRP7-associated mRNAs, resulting in a reduction in translation of defense-related proteins (Nicaise et al., 2013). It has also been shown that Arabidopsis GRP7 regulates alternative splicing of pre-mRNAs and directly binds to pre-mRNAs, modulating alternative splicing (Streitner et al., 2012). All of these observations made GRP7 a prime candidate for further analysis with regard to its role in exRNA production and/or accumulation.

To confirm that GRP7 is secreted into the apoplast, we performed immunoblots on protein isolated from the P40 and P100-P40 fractions of an Arabidopsis line expressing GRP7-GFP expressed under its native promoter (Figure 8). These analyses revealed that GRP7 is mostly detected in the P100-P40 fraction, and therefore likely is not located inside EVs, which mostly pellet in the P40 fraction. To confirm that GRP7 is located outside EVs, we performed a protease protection assay. GRP7 was degraded in the absence of detergent, indicating that it is located outside EVs (Figure 8B).

**Figure 8.**
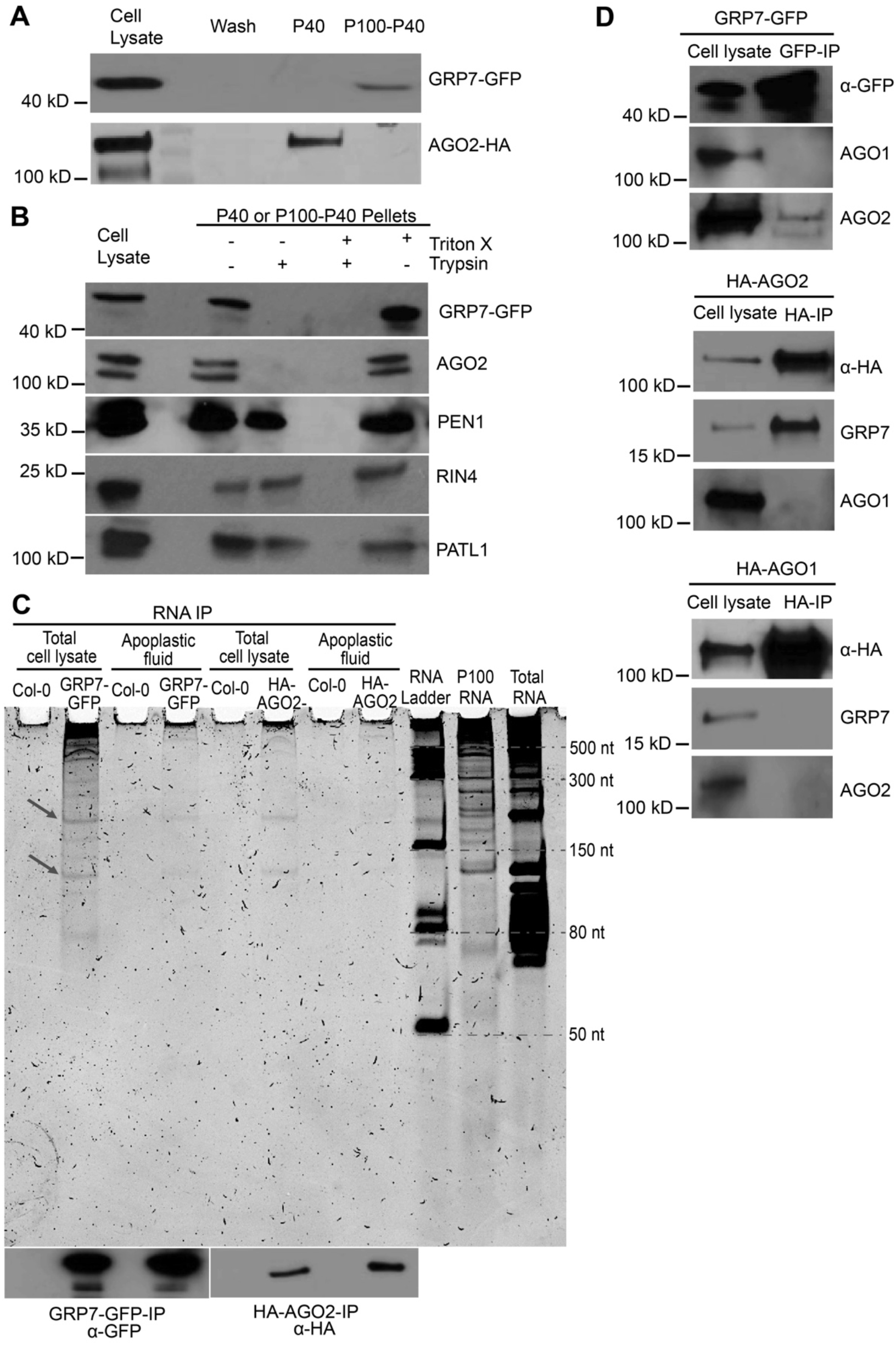
GRP7 and AGO2 Are Secreted to the Apoplast Independent of EVs and Bind to lncRNAs. **(A)** GRP7 and AGO2 are present in the apoplast. Apoplastic fluid was isolated from HA-tagged AGO2 and GFP-tagged GRP7 transgenic Arabidopsis and pelleted at 40,000g (P40) followed by another round of centrifugation at 100,000g (P100-P40). Apoplastic HA-AGO2 mostly pelleted at 40,000g, whereas GRP7-GFP mostly pelleted at 100,000g (P100-P40). Wash; 40 μL of apoplastic wash prior to ultracentrifugation. **(B)** GRP7 and AGO2 are located outside EVs. GRP7- and AGO2-containing pellets were treated with trypsin with or without detergent. GRP7 and AGO2 were eliminated even in the absence of detergent, while known EV cargo proteins PEN1, RIN4 and PATL1 were not. **(C)** GRP7 and AGO2 bind to lncRNAs. RNAs isolated from GRP7-GFP-RNAIP and HA-AGO2-RNAIP were separated by size in a 15% denaturing polyacrylamide gel. Non-transgenic wild-type Arabidopsis was used as a negative control. **(D)** GRP7 and AGO2 co-immunoprecipitate. Whole cell lysates from transgenic plants expressing GRP7-GFP under a GRP7 native promoter, or HA-AGO1, or HA-AGO2 under their respective promoters, were immunoprecipitated with anti-GFP or anti-HA beads. Untagged AGO1, AGO2 and GRP7 proteins were detected using antibodies raised to those proteins. AGO2 coimmunoprecipitated with GRP7-GFP, while AGO1 did not. Reciprocally, anti-HA immunoprecipitation of cell lysates from HA tagged AGO1 and AGO2 plants showed interaction between AGO2 and GRP7, but not between AGO1 and GRP7.

In parallel to our analyses on GRP7, we also assessed whether the sRNA-binding protein ARGONAUTE2 (AGO2) was present in apoplastic wash fluid. AGO2 was chosen from among the ten Arabidopsis AGO proteins because of (1) a known role in plant-pathogen interactions (Harvey et al., 2011), and (2) the availability of a high-quality commercial antibody from Agrisera. Our analyses revealed that, like GRP7, AGO2 is also present in the apoplast, with the majority of it located outside EVs (Figures 8A and 8B).

### GRP7 and AGO2 Associate with lncRNAs in the Apoplast

To investigate the RNAs associated with GRP7 and AGO2 in the apoplast, we performed RNA immunoprecipitation on whole cell lysate and P100 fractions. RNAs were separated by size using polyacrylamide gels, followed by staining with SYBR® Gold to detect nucleic acids. These analyses indicated that Arabidopsis GRP7-GFP, expressed under the native GRP7 promoter, binds to various size RNAs ranging from 50 nt to more than 500 nt in both whole cell lysate and in the P100 fraction (Figure 8C). Similarly, immunoprecipitation of Arabidopsis HA-AGO2 expressed under its native promoter revealed that Arabidopsis AGO2 binds to lncRNAs in both cell lysate and extracellular spaces of plant cells (Figure 8C). In mammalian systems, it has been reported that AGO proteins can bind to lncRNAs through AGO/miRNA complexes (Tarallo et al., 2017). Recently, mammalian AGO proteins have been shown to bind to circRNAs and may function in loading circRNAs into the extracellular matrix (Hansen et al., 2011; Chen et al., 2019; Xu et al., 2020a). The interaction between circRNAs and AGO might be mediated by miRNAs or through interaction with another RNA-binding protein that binds to circRNAs (Hansen et al., 2011; Chen et al., 2019; Zang et al., 2020).

Notably, the RNA banding pattern from AGO2-RNA-IP and GRP7-RNA-IP appears to be similar in polyacrylamide gels (Figure 8C), suggesting that GRP7 and AGO2 may be part of the same RNA-protein complex. We thus performed a co-immunoprecipitation analysis from whole leaf extracts and observed a strong interaction between AGO2 and GRP7 (Figure 8D). Whether the interaction between AGO2 and GRP7 is direct or through binding to the same RNAs is not yet known. Notably, we could not detect an interaction between GRP7 and AGO1, indicating that the GRP7-AGO interaction is specific to AGO2 (Figure 8D).

### Mutation of *AGO2* or *GRP7* Alters Apoplastic circRNA Content

To investigate whether AGO2 and/or GRP7 exert a specific effect on circRNA secretion or stability in the apoplast, we performed RNA-seq analyses on exRNAs from *grp7* and *ago2* mutants following RNase R treatment. These analyses revealed a marked reduction in total circRNAs identified in each mutant (Figure 6B), suggesting that these proteins contribute to circRNA secretion or stabilization.

## DISCUSSION

Prior to the work presented above, it had been unclear whether siRNAs and miRNAs found in the apoplast of plant leaves are primarily packaged inside EVs or are exported via an alternative pathway. In our previous work, we had shown that removal of EVs from apoplastic wash fluid does not deplete the fluid of most siRNAs, suggesting that most siRNAs are located outside EVs (Baldrich et al., 2019). However, (Cai et al., 2018a) reported that siRNAs co-pellet with plant EVs and are resistant to degradation by micrococcal nuclease. Based on these observations, it was concluded that these siRNAs were packaged inside EVs. To address these seemingly contradictory results, we treated EV pellets with protease plus RNase A, which is expected to eliminate sRNAs located outside EVs, but not those located inside EVs. The majority of small RNAs in the size classes of 21, 22 and 24 nt were eliminated (Figure 1A), which indicates that most siRNAs and miRNAs are not located inside EVs but are located outside EVs and are protected from nucleases by RNA-binding proteins. This finding is consistent with recent work in mammalian systems, which has shown that many sRNAs that co-purify with EVs can be digested with protease plus RNase treatment (Shurtleff et al., 2017; Jeppesen et al., 2019) and are thus likely located outside EVs. This finding also suggests that EVs may not play a direct role in translocating sRNAs into other organisms such as fungal pathogens. Instead, it appears that sRNA-protein complexes located outside EVs could be the primary mediators of interkingdom RNA silencing.

Although our data indicate that the majority of sRNAs are located outside EVs, it is important to note that many sRNAs co-pellet with EVs during differential ultracentrifugation. This could be because the sRNAs are bound to protein complexes of a size similar to that of EVs and/or they could be associated with the surface of EVs. EVs have a relatively high surface area in comparison to their volume, which can promote interactions between EVs and other extracellular molecules (Janas et al., 2015; Buzás et al., 2018). A tight association between sRNAs and EV surface proteins could potentially protect sRNAs from degradation by nucleases.

In addition to sRNAs, our analyses of apoplastic RNAs revealed that plants secret lncRNAs into the extracellular space. Although it has been reported that some extracellular lncRNAs are located inside mammalian EVs (Takahashi et al., 2014; Chen et al., 2016; Zheng et al., 2018; Dai et al., 2020) our data indicate that extracellular lncRNAs produced by plants are located outside EVs and are associated with RNA-binding proteins. As with sRNAs, we found it was necessary to treat apoplastic pellets with protease prior to RNase A to determine whether lncRNAs were inside or our outside EVs, as treatment with RNase A alone had very little effect (Figure 5A).

In mammalian systems, lncRNAs have been shown to regulate multiple biological processes, including gene transcription (Luo et al., 2016), translation (Hu et al., 2018; Jia et al., 2019) and epigenetic modifications (Neumann et al., 2018), as well as cell-to-cell communication (Wei and Wang, 2015; Cai et al., 2018b; Zhu et al., 2021). lncRNAs have also been shown to contribute to antiviral innate immune responses in mammalian systems (Ouyang et al., 2016; Liu et al., 2020). Similarly, lncRNAs in plants have also been shown to modulate gene expression, epigenetic regulation and response to stresses (Di et al., 2014; Wang et al., 2018; Hamid et al., 2020; Moison et al., 2021). However, the presence of lncRNAs in the extracellular space of plant cells and their roles in cell-to-cell communication or immune responses have not been investigated yet. Whether plant extracellular lncRNAs can be taken up by pathogen cells is unknown, but the ability of fungi to take up long single-stranded and double-stranded RNAs in a petri dish suggests that this is likely (Qiao et al., 2021). If so, it will be interesting to assess whether these RNAs can impact gene expression in fungi and other plant-associated organisms.

A subclass of lncRNAs of particular interest is circRNAs, as these have previously been shown to be induced by pathogen infection in plants, and circRNAs appear to contribute to immunity (Fan et al., 2020). Our sequencing data revealed that Arabidopsis exRNA contains thousands of circRNAs. At the same time, no intact full-length mRNAs were identified, indicating that circRNAs are preferentially secreted or are more stable in the apoplast than linear mRNAs. This finding is similar to that reported for cultured human cells, in which circRNAs were found to co-purify with EVs and to be highly enriched relative to their matching linear RNAs found in cell lysates (Lasda and Parker, 2016).

Extracellular circRNAs in mammals have been suggested to contribute to cell-to-cell communication (Lasda and Parker, 2016). One likely function of mammalian extracellular circRNAs is as a sponge for sequestering miRNAs (Hansen et al., 2013). Whether plant circRNAs play a similar role in the apoplast is not yet known, but it is tempting to speculate that they could function as target mimics for small RNAs secreted by pathogens. Pathogens have been reported to deliver sRNAs into plant cells to suppress immunity and enhance susceptibility (Weiberg et al., 2013; Wang et al., 2017; Dunker et al., 2020); thus, having a collection of sponges in the apoplast to soak up sRNAs secreted by pathogens before they can reach their targets inside the host cell could be quite useful.

The discovery that plants accumulate lncRNAs in their extracellular spaces raised the fundamental question of how this RNA is secreted. We found that the RNA-binding proteins AGO2 and GRP7 also accumulate in the apoplast and are bound to lncRNAs. Elimination of these proteins altered the RNA content of the apoplast, which indicates a possible function of AGO2 and GRP7 in the secretion of RNA into the apoplast or stabilizing RNAs once there. Notably, GRP7 belongs to the same family of RNA-binding proteins as human HNRNPA2B1, which has been shown to mediate sorting of specific miRNAs into EVs (Villarroya-Beltri et al., 2013), and to bind to m^6^A-modified RNA (Alarcón et al., 2015), which suggests that GRP7 could be fulfilling similar roles in plants. Consistent with this hypothesis, we found that plant exRNAs are highly enriched in m^6^A modifications. Whether m^6^A modification plays a role in the secretion of exRNAs into the apoplast or contributes to their stability requires further investigation.

## METHODS

### Plant Materials and Growth Conditions

The *Arabidopsis thaliana grp7* mutant (SALK_039556.21.25.x) was obtained from the Arabidopsis Biological Resource Center at Ohio State. The Arabidopsis *ago2-1* mutant was obtained from James Carrington at the Donald Danforth Plant Science Center. The Arabidopsis HA-AGO2 transgenic line was also obtained from Dr. Carrington. It expresses HA-AGO2 under the native *AGO2* promoter in an *ago2-1* mutant background (Montgomery et al., 2008). The GRP7-GFP transgenic line was obtained from Dr. Dorothee Staiger at Bielefeld University. This line expresses GRP7-GFP under control of the native *GRP7* promoter and the *GRP7* 5’UTR, intron and 3’UTR in a *grp7–1* mutant background (Köster et al., 2014). Seeds were germinated on 0.5X Murashige and Skoog medium containing 1% agar. To induce synchronous germination, petri dishes containing the seeds were stored at 4°C for 2 days and then moved to short-day conditions illuminated using GE HI-LUMEN XL Starcoat 32 watt fluorescent bulbs (a 50:50 mixture of 3,500K and 5,000K spectrum bulbs; 9 hour days, 22°C, 150 μEm^−2^s^−1^). After 10 days, the seedlings were transferred to Pro-Mix FLX potting mix supplemented with Osmocote slow-release fertilizer (14-14-14). Seedlings were grown under a clear plastic dome for the first week following transfer.

### Isolation of EVs and Other Apoplastic Particles

Apoplastic wash fluid was isolated from 6-week-old Arabidopsis plants as described in Rutter and Innes (2017). Briefly, Arabidopsis rosettes were vacuum-infiltrated with vesicle isolation buffer (VIB), pH 6.0, containing 20 mM 2-(N-morpholino) ethanesulfonic acid (MES), 2 mM CaCl_2_, and 0.01 M NaCl as described previously (Rutter et al., 2017). After vacuum-infiltration, the excess buffer was removed from leaf surfaces by blotting rosettes with Kimwipes®. To recover apoplastic fluid from infiltrated leaves, rosettes were placed inside needleless, 30-mL syringes (two rosettes per syringe). Syringes were placed inside 50-mL tubes and centrifuged for 20 min at 700g with slow acceleration (4°C, JA-14 rotor, Avanti J-20 XP Centrifuge; Beckman Coulter). The apoplastic wash fluid was then filtered through a 0.22 μm membrane and centrifuged at 10,000g for 30 minutes to remove any remaining large particles. The supernatant was transferred into new centrifuge tubes and centrifuged at 40,000g (P40) or 100,000g (P100) for one hour (4°C, TLA100.3, Optima TLX Ultracentrifuge; Beckman Coulter) to pellet EVs and other particles as noted in figure legends. The pellet was washed and re-pelleted at 40,000g or 100,000g at 4°C using a TLA100.3 rotor, Optima TLX Ultracentrifuge (Beckman Coulter). The pellets were re-suspended in 100 μL of cold and filtered VIB (0.22 μm) and either used immediately, or stored at −80°C until further use.

### RNA Purification

Total leaf RNA was isolated from 100 mg of fresh or frozen leaf tissue using TRIzol Reagent (Thermo Fisher Scientific). Briefly, to isolate RNA, leaf tissue was frozen in liquid nitrogen and ground into powder using a mortar and pestle. One mL of TRIzol Reagent (Thermo Fischer Scientific, Waltham, MA) was added to the ground leaf tissue and mixed vigorously by vortexing. The leaf and TRIzol mixture was then shaken at room temperature for 10 minutes, followed by the addition of 200 μL of chloroform. This mixture was then vortexed for 30 seconds and then centrifuged at 12,000g for 15 minutes. The aqueous phase was removed and mixed with one volume of cold isopropanol to precipitate the RNA. RNA pellets were washed using 80% cold ethanol. To isolate RNA from P40 and P100 pellets, 1 mL of TRIzol was added to 100 μL of resuspended pellet, followed by the same procedure as used for leaf RNA isolation. RNA pellets were re-suspended in 10 to 12 μL of ultrapure DNase/RNase-free water (Invitrogen) and stored at −80°C. RNA quality and quantity was assessed using either a ThermoFisher NanoDrop One spectrophotometer, or an Agilent 2200 Tape Station.

### Trypsin and RNase A Treatments

To assess whether RNAs were located inside or outside EVs, we performed RNase protection assays as follows. P40 pellets were treated with 1 μg/mL trypsin (Promega) in the presence or absence of 1% (v/v) Triton X-100 (EMD-Millipore) in 15 mM Tris-HCl (80 ul final volume). Samples were incubated at 37°C for one hour followed by adding 1.5 μg/mL trypsin inhibitor (Worthington Biochemical. Corp) to inactivate trypsin. For the samples with RNase treatment, RNase A (Qiagen; diluted in 15 mM NaCl, 10 mM Tris-HCl pH 7.5) was added to the mixture to a final concentration of 5 μg/mL (100 μL final volume) and the sample was incubated at room temperature for 30 minutes. Immediately after RNase A treatment, RNA was isolated using 1 mL of TRIzol as described above. To inhibit RNase A activity, a mixture of 10 μg/mL RNase Inhibitor, Murine (APExBIO) and 40 unit/mL of RNase Out (Invitrogen) was added to the RNAs and stored at −80°C until library preparation.

### RNA-Immunoprecipitation (RNA-IP)

To isolate RNAs associated with GRP7-GFP and AGO2-HA from whole leaves and from apoplastic fluid of transgenic Arabidopsis plants we performed RNA-IP. For leaves, we used one gram of fresh or frozen leaf tissue, which was frozen under liquid nitrogen and ground with a mortar and pestle. Leaf powder was mixed with 5 mL of cold IP buffer (0.05 M Tris-Hcl, pH 7.4, 0.1 M KCl, 2.5 mM MgCl_2_, 0.1 % NP-40, 1% Triton X-100 and 50 U/mL RNase Out), incubated on ice for 10 minutes, and then transferred to a 15 mL polypropylene screw-cap centrifuge tube. The tube was then centrifuged for 10 minutes at 12,000g and the supernatant was filtered through a 0.45 μm membrane. The filtered supernatant was then incubated for one hour at 4°C with 50 μL of anti-GFP agarose beads (Chromotek) to pull down GRP7-GFP and with 50 μL of anti-HA agarose beads (ThermoFisher) to pull down AGO2-HA. Beads were then pelleted by centrifugation at 1000g for 2 minutes at 4°C and washed at least 6 times with 5 mL of cold IP buffer for 5 minutes at 4°C for each washing step. Finally, beads were washed two times with 1.5 mL of cold IP buffer followed by a final wash with 1 mL of ultrapure RNase-free/DNase-free water, and pelleted by centrifugation at 538g for 1 minute. To immunoprecipitate GRP7-GFP and AGO2-HA from apoplastic fluid, P100 pellets were re-suspended in 2 mL of cold IP buffer and proteins immunoprecipitated as described for whole leaf extracts.

To isolate RNA, beads were incubated with proteinase K at a final concentration of 1.5 μg/μL in 100 μL of PK buffer (0.1 M Tris-HCl, pH 7.4, 0.01 M EDTA, pH 8.0, 300 mM NaCl and 2% SDS) for one hour at 55°C with intermittent shaking (every 3 minutes for 15 seconds). Beads were pelleted by centrifugation at 538g for 1 minute and RNA isolation was performed using TRIzol reagent as described above.

### Polyacrylamide Gel Preparation and Electrophoresis

RNA samples were analyzed using denaturing polyacrylamide gel electrophoresis. Gels containing 15% polyacrylamide and 7 M urea were prepared using IBI InstaPAGE 40% acrylamide solution (37.5:1). RNA samples were denatured at 65°C in denaturing buffer (0.25 M EDTA (pH 8.0), 8 M Urea, 0.2 mg/mL bromophenol blue, 0.02 mg/mL xylene cyanol) and resolved on 0.5 x Tris-Boric Acid EDTA (0.5 x TBE; 0.065 mM Tris (pH 7.6), 21 mM boric acid, 1.25 mM EDTA)-15% polyacrylamide urea gels. For size standards, we used New England Biolabs Low Range ssRNA Ladder (catalog number N0364S) and Takara 14-30 ssRNA Ladder Marker (catalog number 3416). SYBR® Gold Nucleic Acid Gel Stain (ThermoFisher) was used to stain gels for 30 minutes before UV transillumination. Gel images were acquired using a Bio-Rad ChemiDoc-MP imaging system.

### RNA Dot Blots Using Anti-m6A Antibodies

RNA was isolated from leaf or apoplastic P40 and P100 fractions using TRIzol as described above and the RNA concentration was measured using a ThermoFisher NanoDrop One spectrophotometer. For all samples, equal amounts of RNA were prepared in equal volumes (6 μL) using UltraPure DNase/RNase-free distilled water (Invitrogen™). RNA samples were denatured at 95°C for 3 minutes and placed on ice immediately to prevent the formation of secondary structures. A piece of Hybond-N+ membrane (Amersham Pharmacia Biotech) was prepared and RNA samples were applied directly to the Hybond-N+ membrane using a micropipettor. To prevent the spread of RNA on the membrane, 2 μL of RNA solution was applied at a time, allowing the membrane to dry for three minutes before applying the next 2 μL drop to the same spot, until a total of 6 μL of RNA sample was applied. To crosslink the spotted RNAs to the membrane, an UVC-508 Ultraviolet Cross-linker (Ultra-Lum) was used to irradiate the membrane twice at 1200 microjoules [x100] for 30 seconds. The membrane was then washed in clean RNase-free 1x PBS buffer (1x PBS; 2.7 mM KCl, 8 mM Na2HPO4, 2 mM KH2PO4 and 137 mM NaCl, pH 7.4) and blocked in 5% non-fat milk in 1x PBS containing 0.02% Tween-20 for one hour at room temperature. The membrane was then incubated overnight with anti-m^6^A antibody (Abcam catalog number ab151230) at a 1:250 dilution in 5% non-fat milk in 1x PBS containing 0.02% Tween-20. The membrane was washed in 1x PBS containing 0.02% Tween-20 three times and incubated with horseradish peroxidase-labeled goat anti-rabbit antibody (Abcam catalog numbe ab205718) at a 1:5000 dilution for 1 hour (Lisha et al., 2017). After final wash in1x PBS contain 0.02% Tween-20, m^6^A modified RNAs were visualized using Immune-Star Reagent (Bio-Rad) and imaged using X-ray film.

### Preparation of Circular RNA Samples

To investigate the presence of circRNAs, RNA was isolated from 100 μL of P100 pellet using a PicoPure RNA isolation kit (ThermoFisher). The RNA (1-3 μg) was then treated with 5 units of RNase R (Lucigen. RNR07250) for one hour at 37°C. To visualize circRNAs, the RNA samples were resolved on TBE-15% polyacrylamide urea gels and stained with SYBR® Gold. To prepare RNA libraries for sequencing, it was necessary to remove RNase R from the RNA samples. This was accomplished by re-purifying the RNase R-treated RNA samples using a PicoPure RNA isolation column.

### Preparation of sRNA-seq and RNA-seq Libraries

sRNA libraries were constructed using the RealSeq-AC kit (no. 500-00048; RealSeq Biosciences) following the manufacturer’s recommendations. To capture all types of small RNAs, we used 1 μg of RNA as starting material. Except for RNase-R treated samples, all RNA-seq libraries were generated using the NEBNext® Ultra™ II Directional RNA Library Prep Kit for Illumina® (catalog number E7765; New England Biolabs) using 500 ng of total RNA as starting material. rRNA removal was accomplished using the RiboMinus™ Plant Kit for RNA-Seq (catalog number A1083808, ThermoFisherScientific) and Poly(A) RNA purification was attempted using the NEBNext® Poly(A) magnetic isolation module (catalog number E7490, New England Biolabs). For sequencing of RNase R-treated samples, RNA-seq libraries were prepared using an Illumina TruSeq Stranded mRNA Library Prep kit (catalog number 20020594; Illumina) following the manufacture’s protocol, but skipping the poly(A) enrichment step. All libraries were sequenced on an Illumina NextSeq 550 instrument with paired-end 75-bp reads, except for the RNase R-treated samples, which were sequenced using paired-end 300-bp reads. Sequencing was performed at the Center for Genomics and Bioinformatics at Indiana University, Bloomington (IN, USA).

### Data Analysis

sRNA sequencing libraries were trimmed of adaptors using the software Cutadapt v1.16 (Martin, 2011) with a minimum insert size of 10 nt and a maximum of 34 nt. Sequence quality was assessed using FastQC (http://www.bioinformatics.babraham.ac.uk/projects/fastqc/). Clean reads were aligned to the Arabidopsis genome (TAIR version 10), and all subsequent analyses were performed using the software Bowtie2 (Langmead and Salzberg, 2012). For miRNA analyses, the latest version of miRBase (v22; (Kozomara and Griffiths-Jones, 2014) was used. RNA-seq libraries were also trimmed of adaptors using Cutadapt v1.16 (Martin, 2011) and sequence quality assessed using FastQC. Clean reads were aligned to the Arabidopsis genome (TAIR version 10), using HISAT2 version 2.2.1 (Kim et al., 2019). To identify circRNAs, mapping was performed using the Arabidopsis data on PlantcircBase v5.0 (http://ibi.zju.edu.cn/plantcircbase/) (Chu et al., 2017). We only considered reads mapping concordantly and exclusively to the junction part of the circular RNA. Differential accumulation analyses were performed using DEseq2 with default parameters, using not-normalized reads as input (Love et al., 2014). Graphical representations were generated using the software ggplot2 (Wickham, 2009) in the R statistical environment.

---

**Supplemental Figure 1.**
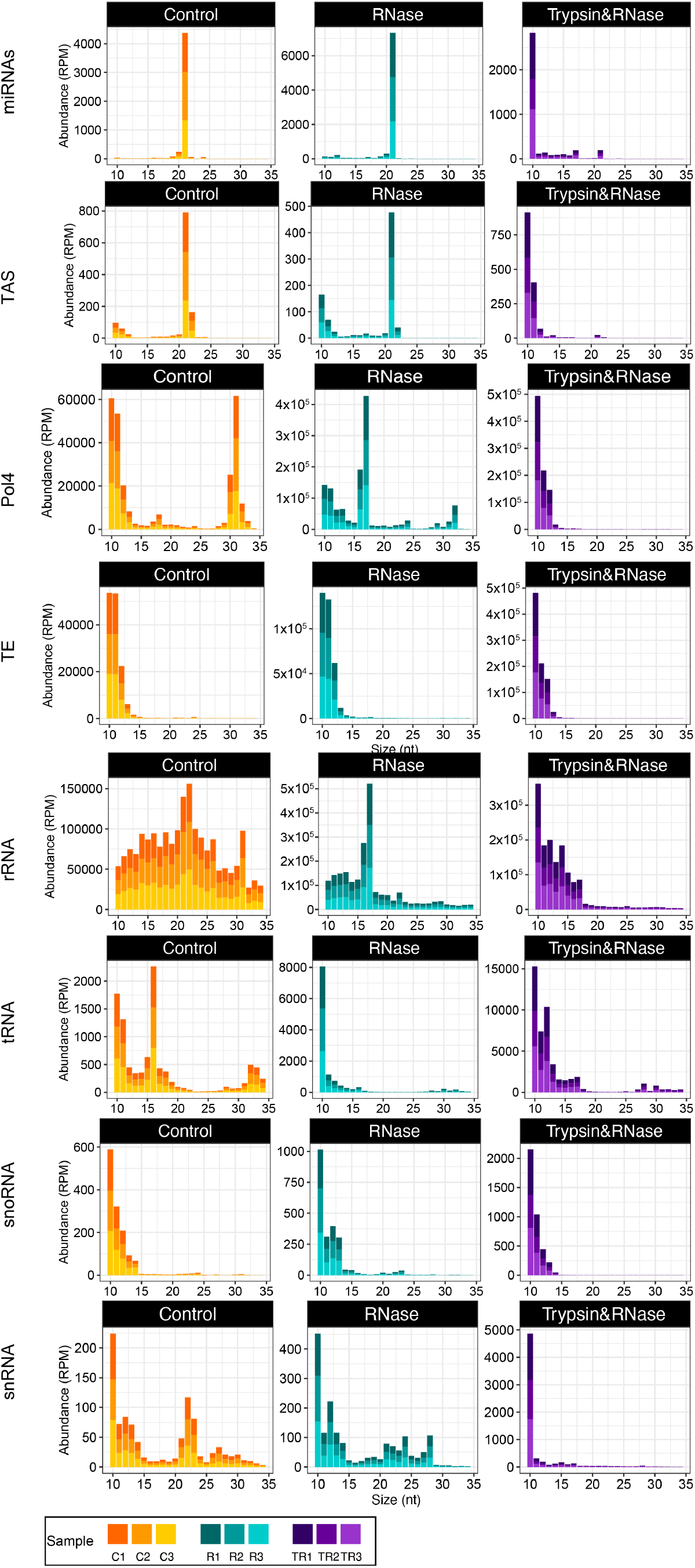
Apoplastic miRNAs and trans-acting siRNAs are mostly located outside EVs and are protected by proteins. Graphs indicate the size distributions of P40 sRNAs mapping to the indicated sources. The abundance of each size class was calculated for each P40 treatment: control (C), RNase A only (R), and trypsin plus RNase A (TR). The x axis indicates the sRNA size and the y axis indicates its abundance in reads per million (RPM). Data from three independent biological replicates are stack together in a single bar plot and color coded as shown in the legend.

**Supplemental Figure 2.**
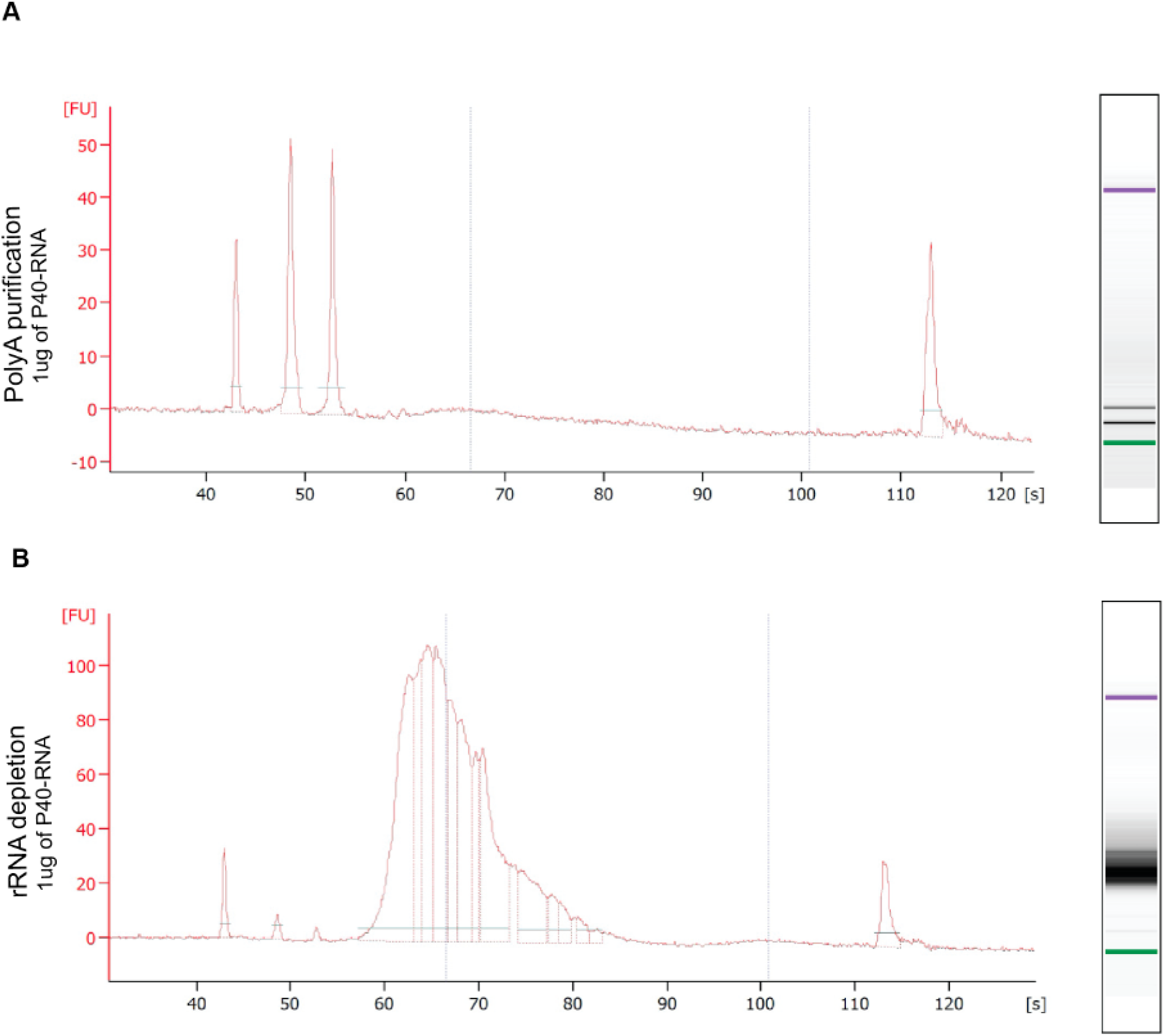
P40 RNA Appears to Lack Poly-Adenylated RNA. RNA-seq libraries were prepared from P40 RNA using two different methods. **(A)** Method 1 employed a poly(A) enrichment step to specifically copy poly-adenylated mRNAs. Analysis of the sizes of the inserts in the resulting library using an Agilent Tape Station revealed that most products lacked an insert, indicating a lack of full-length mRNAs in the P40 fraction. **(B)** The second method used a ribosomal RNA depletion step, but no poly(A) enrichment step. This library produced products with the expected size range of inserts (note broad peak between 60 and 80 seconds).

## Accession Numbers

The data discussed in this publication have been deposited in NCBI’s Gene Expression Omnibus (Edgar et al., 2002) and are accessible through GEO Series accession numbers GSE183867 (https://www.ncbi.nlm.nih.gov/geo/query/acc.cgi?acc=GSE183867) and GSE185133 (https://www.ncbi.nlm.nih.gov/geo/query/acc.cgi?acc=GSE185133). The accession numbers for Arabidopsis proteins discussed in this work are AT1G48410 (AGO1), AT1G31280 (AGO2), AT2G21660 (GRP7), AT1G72150 (PATL1), AT3G11820 (PEN1), AT1G59870 (PEN3), and AT3G25070 (RIN4).

## Supplemental Data

**Supplemental Figure 1**. Apoplastic miRNAs and trans-acting siRNAs are mostly located outside EVs and are protected by proteins.

**Supplemental Figure 2**. Apoplastic RNA appears to lack poly-adenylated RNA.

## ACKNOWLEDGMENTS

We thank Dorothee Staiger at the University of Bielefeld for providing seed of *GRP7-GFP* transgenic Arabidopsis and Jim Carrington for providing seed of *ago2-1* mutant and HA-AGO2 transgenic Arabidopsis. We also thank the Arabidopsis Biological Resource Center at Ohio State for proving seed of an Arabidopsis *grp7* mutant, and the Indiana University Physical Biochemistry Instrumentation Facility for access to ultracentrifuges and nanoparticle tracking equipment. We also thank the IU Center for Genomics and Bioinformatics for assistance with the generation and analysis of sRNA-seq and RNA-seq data. This work was supported by three grants from the United States National Science Foundation, IOS-1645745 and IOS-1842685 to RWI and IOS-1842698 to BCM.

## AUTHOR CONTRIBUTIONS

H.Z.K., P.B., B.D.R., R.W.I. and B.C.M. designed the research; H.Z.K., P.B., B.D.R., K.Z. and L.B. performed the research; P.B. analyzed the sRNA-seq and RNA-seq data; H.Z.K., P.B., and R.W.I. wrote the article; all authors read and commented on the article.

